# A detailed characterisation of drug resistance during darunavir/ritonavir (DRV/r) monotherapy highlights a high barrier to the emergence of resistance mutations in protease but identifies alternative pathways of resistance

**DOI:** 10.1101/2023.10.15.562382

**Authors:** Adam Abdullahi, Ana Garcia Diaz, Olga Mafotsing Fopoussi, Apostolos Beloukas, Victoire Fokom Defo, Charles Kouanfack, Judith Torimiro, Anna Maria Geretti

## Abstract

**Background:** Maintenance monotherapy with DRV/r has yielded variable outcomes and is not recommended. Trial samples offer valuable opportunities for detailed studies. We analysed samples from a 48-week trial in Cameroon to obtain a detailed characterisation of drug resistance.

**Methods:** Following failure of NNRTI-based therapy and virological suppression on PI-based therapy, participants were assigned to receive either DRV/r (n=81) or TDF/3TC + LPV/r (n=39). PBMC from study entry underwent bulk protease and RT sequencing. Plasma collected at virological rebound (confirmed or last available HIV-1 RNA ≥60 copies/ml) underwent ultradeep protease and RT sequencing and bulk gag-protease sequencing. A site-directed mutant with T375A (p2/p7) was phenotypically characterised using a single-cycle assay.

**Results:** HIV-1 DNA analysis revealed NRTI and NNRTI resistance-associated mutations (RAMs) in 52/90 (57.8%) and 53/90 (58.9%) samples, respectively. In rebound HIV-1 RNA (DRV/r n=21; LPV/r n=2), prevalence was 9/23 (39.1%) and 10/23 (43.5%), respectively, with most RAMs occurring at frequencies ≥15%. No DRV RAMs were found. Paired HIV-1 DNA and RNA sequences showed partial consistency in resistance patterns. Among 8 participants experiencing virological rebound on DRV/r (n=12 samples), all showed gag mutations associated with PI-exposure, including T375N, T375A (p2/p7), K436R (p7/p1), and mutations in p17, p24, p2 and p6. T375A conferred 10-fold DRV resistance and increased replication capacity.

**Conclusions:** The study highlights the high resistance barrier of DRV/r while identifying alternative pathways of DRV resistance through gag substitutions. During virological suppression, resistance patterns in HIV-1 DNA reflect treatment history, but due to technical and biological considerations, cautious interpretation is warranted.

## Introduction

HIV drug resistant variants acquired at the time of infection or enriched under selective drug pressure can integrate into the DNA of memory CD4 T-cells as provirus and become part of the HIV DNA archive within the host cell.^1-2^ Archived resistant variants can re-emerge if virus production resumes, thus potentially retaining life-long clinical significance. While the available evidence is not completely consistent, studies have demonstrated that detecting drug resistance-associated mutations (RAMs) in the HIV-1 DNA of virologically suppressed patients predicts virological outcomes when switching to a different treatment regimen.^2,3^ Interestingly, some studies have produced surprising finding. We previously reported that detecting archived RAMs during suppressive antiretroviral therapy (ART) with of 2 nucleos(t)ide reverse transcriptase inhibitors (NRTIs) plus a ritonavir-boosted protease inhibitor (PI/r) was associated with a reduced likelihood of virological rebound after switching to darunavir/ritonavir (DRV/r) monotherapy.^4^

Maintenance monotherapy with DRV/r has been studied in Western Europe among virologically suppressed patients without a previous treatment failure.^5^ A multi-centre study conducted in Burkina Faso, Cameroon and Senegal investigated DRV/r monotherapy among 50 patients who had achieved virological suppression on PI/r-based triple ART after failure of 2 NRTIs plus a non-nucleoside reverse transcriptase inhibitor (NNRTI).^6^ These studies uniformly reported an increased risk of viremia when switching to DRV/r monotherapy, and also consistently demonstrated a low risk of treatment emergent DRV RAMs.

We conducted a randomised clinical trial In Cameroon comparing DRV/r maintenance monotherapy to standard of care (SOC) triple ART with 2 NRTIs plus lopinavir/ritonavir (LPV/r) in adults living with HIV. Similar to the study from Burkina Faso, Cameroon and Senegal,^6^ participants were virologically suppressed on PI/r-based triple ART having experienced virological failure of first-line NNRTI-based therapy. We previously reported the virological outcomes of the DRV/r arm in relation to the detection of RAMs in cellular HIV-1 DNA at study entry.^4^ The aim of this analysis was to take advantage of unique trial samples and conduct a detailed characterisation of the virological outcomes of the entire trial population. We performed a sensitive assessment of treatment-emergent resistance using ultradeep sequencing (UDS), included the evaluation of gag mutations, and compared the resistance patterns detected in cellular HIV-1 DNA at study entry, when participants showed virological suppression, to those observed in plasma HIV-1 RNA during subsequent virological rebound.

## Methods

### Study population

MANET (Monotherapy in Africa, New Evaluations of Treatment) was a randomised, open-label trial based at Hôpital Central Yaoundé in Cameroon (NCT02155101). Between August 2014 and July 2015, 120 adults receiving virologically suppressive ART with 2 NRTIs plus a PI/r (typically LPV/r) were randomised to either DRV/r monotherapy (800/100mg once daily) for 48 weeks (n=81) or standard of care (SOC) with tenofovir disoproxil fumarate co-formulated with lamivudine (TDF/3TC) plus LPV/r for 24 weeks (n=39). Eligibility criteria comprised having received 2 NRTIs plus a PI/r for ≥12 weeks, CD4 count >100 cells/mm^3^, plasma HIV-1 RNA <60 copies/ml in two screening measurements taken 4-12 weeks apart, negative hepatitis B surface antigen (HBsAg) and absence of significant disease or laboratory abnormalities. Pregnancy or planning to become pregnant were exclusion criteria. Following randomisation, participants of both arms attended scheduled study visits at weeks 4, 12 and 24; the DRV/r arm also attended scheduled study visits at weeks 36 and 48. The study was approved by the University of Liverpool Ethics Committee (RETH000605) and the Cameroon National Ethics Committee (2013/07/347) and overseen by an independent trial safety board.

### Statistical analysis

The characteristics of the population at study entry, summarised as categorical and continuous variables, were compared by chi-squared test, Fisher’s exact test or Wilcoxon Mann Whitney test, as appropriate. The trial primary endpoint was the proportion of participants with plasma HIV-1 RNA <400 copies/ml at week 24 (FDA snapshot). Secondary virological endpoints measured through week 24 in both arms and through week 48 in the DRV/r arm comprised proportions with plasma HIV-1 RNA <60 copies/ml and emergence of RAMs in participants with virological rebound, defined as confirmed (or last available) HIV-1 RNA ≥60 copies/ml. Virological failure was defined as confirmed (or last available) HIV-1 RNA ≥400 copies/ml.

### Laboratory tests

At the Centre Pasteur of Cameroon in Yaoundé, safety parameters and CD4 cell counts were measured using freshly collected samples. Plasma was separated from whole venous blood in EDTA within 2 hours of collection and stored at -80°C. HIV-1 RNA was quantified with the Biocentric assay (Bandol, France; lower limit of quantification 60 copies/ml). At the Chantal Biya International Reference Centre for Research on HIV/AIDS Prevention & Management in Yaoundé, peripheral blood mononuclear cells (PBMC) were isolated by Ficoll-Hypaque gradient centrifugation and stored at -80°C. Sanger sequencing of protease (amino acids, aa 1-99) and reverse transcriptase (RT, aa 1-335) was performed as described.^7^ Sample aliquots were shipped frozen to the UK for HIV-1 RNA sequencing (see below) and for the quantification of total HIV-1 DNA in PBMC by real-time PCR as described.^8^

### HIV-1 RNA sequencing

Plasma samples from participants experiencing virological rebound underwent UDS as previously described.^9,10^ Briefly, samples with HIV-1 RNA <10,000 copies/ml were enriched by ultra-centrifugation at 35000 rpm for 20 minutes at 4°C; following extraction with the QIAamp viral RNA kit (Qiagen, UK), a 1300bp amplicon was generated covering protease (aa 1-99) and RT (aa 1-335), purified with Agencourt AMPure XP magnetic beads (Beckman Coulter, UK) and quantified by the Qubit dsDNA High Sensitivity Assay Kit on the Qubit 3.0 fluorometer (Invitrogen, UK). A DNA library was prepared with the Nextera XT DNA Sample Prep Kit (Illumina, USA), followed by sequencing with the Illumina MiSeq Reagent Kit v2. After checking for quality, reads were analysed applying a frequency threshold of 1% and described as low frequency variants (representing 1-14% of the variants in a sample) and high-frequency variants (≥15%). Using plasma samples from participants experiencing virological rebound on DRV/r, after extraction with the QIAamp Viral RNA kit (Qiagen), viral RNA was reverse transcribed using Superscript III One-Step RT PCR Kit with Platinum® Taq High Fidelity enzyme followed by amplification using Platinum® PCR SuperMix High Fidelity (Invitrogen). A 2200bp amplicon spanning gag and protease was generated by nested PCR as detailed in Supplementary Table 1, followed by Sanger sequencing.

**Table 1.**
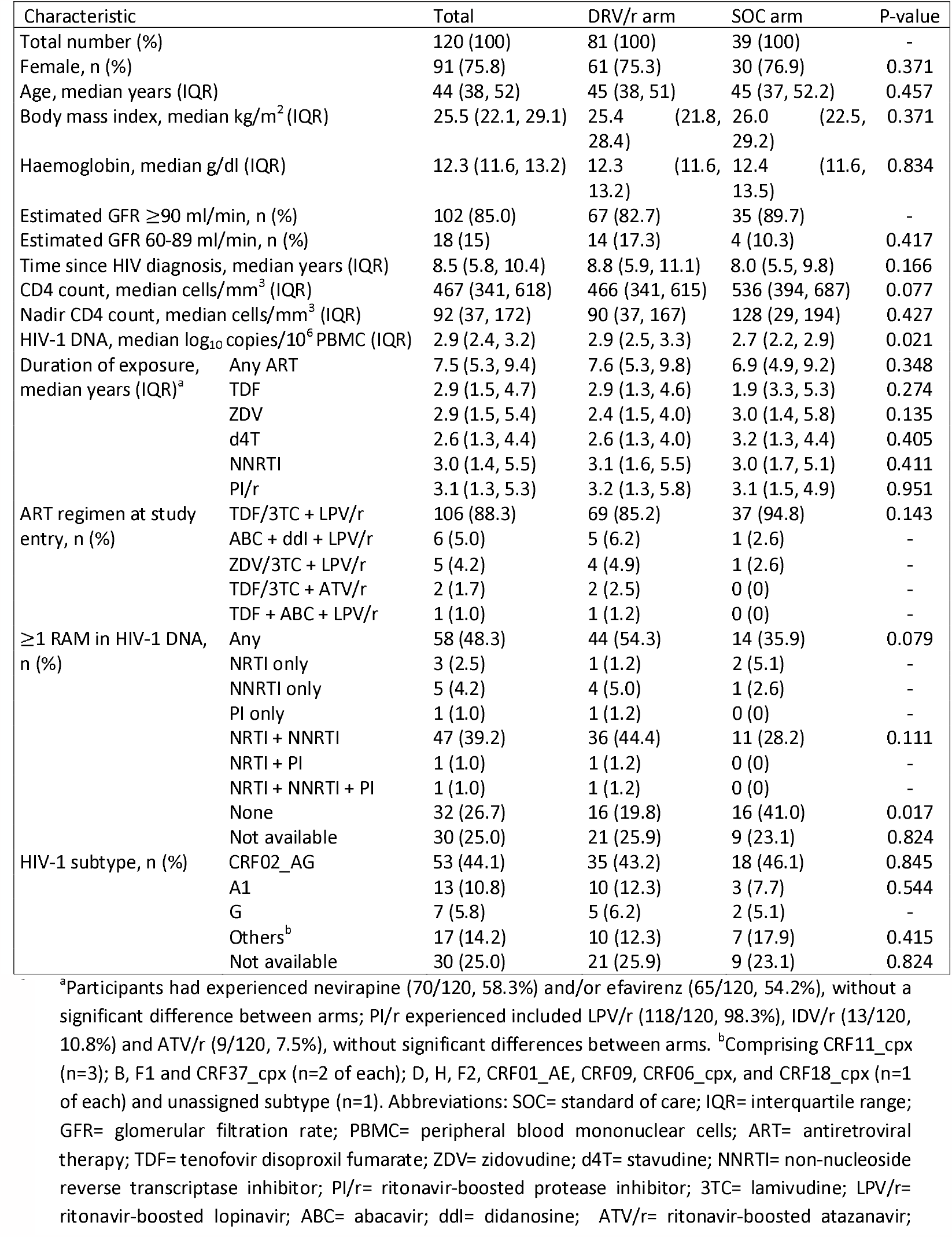
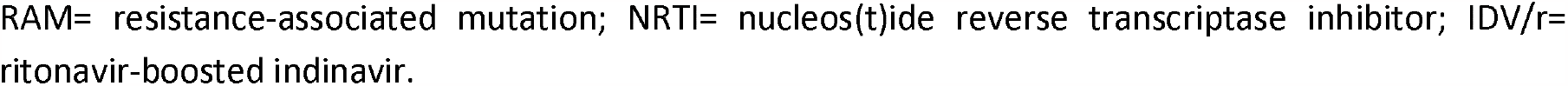
Characteristics of the population at study entry (n=120)

### Sequence analysis

Major RAMs in RT and protease were defined according to the Stanford HIV Drug Resistance Database v9.4.1.^11^ Darunavir RAMs comprised V11I, V32I, L33F, I47V, I50V, I54L/M, T74P, L76V, I84V and L89V. Sequences were analysed for the presence of in-frame stop-codons to indicate defective proviruses and screened for evidence of APOBEC3G (A3G) hypermutation using the Los Alamos Hypermut 2.0 tool.^12^ Phylogenetic analysis was used to investigate linkage between *pol* sequences as described.^13^ Briefly, for each FASTQ and FASTA *pol* sequence generated, 10 reference sequences were downloaded from GENBANK,^14^ duplicate sequences were manually removed, and maximum likelihood phylogenetic trees were constructed using RAxML version 8.^15^ Phylogenies were inferred with Figtree v1.4.4 with 1000 bootstrap replicates.^16^ Phylogenetic analysis was also used to confirm the HIV-1 subtypes. Gag sequences were aligned with the HIV-1 HXB2 reference sequence using MEGA v 6.06.^17^ Mutations were reported according to their association with PI exposure, which was pre-determined by comparing full-length gag and protease sequences from 200 PI-naive and 191 PI-experienced individuals.^18^ Mutations associated with PI exposure were those showing a significantly higher prevalence in the PI-experienced group by Fisher’s exact test with Bonferroni correction, using a p-value threshold of <0.001 for mutations occurring at cleavage sites (p17/p24, p2/p7, p7/p1, p1/p6, p24/p2) and <0.0001 for mutations occurring in other regions of gag. They included 14 cleavage site mutations [2 in p17/p24 (V128I, Y132F), 4 in p2/p7 (S373T, A374S, T375A, T375N), 3 in p7/p1 (A431V, K436R, I437V), and 5 in p1/p6 (L449F, S451T, S451R, R452S, P453T)], and 19 mutations in other regions of gag [10 in p17 (L61I, I94V, K103R, K113Q, K114R, D121G, D121A, T122E, N126S, Q127K), 5 in p24 (T186M, T190I, A210S, E211D, S310T), 3 in p6 (F463L, T469I, P478Q), and 1 in p2 (T371Q)].

### Phenotypic characterisation of the gag cleavage site mutation T375A

T375A was inserted by site directed mutagenesis into the wild-type vector P8.9NSX, which contains the protease and RT sequences of the NL4-3 strain of HIV-1,^19^ using the Quickchange Multi/Site Directed Mutagenesis kit (Strategene, UK). The amplified DNA was enriched by digestion of the parental DNA with DpnI and XL1-blue and supercompetent cells were transformed with the digested DNA. The plasmidic DNA was isolated using QIAPrep Spin Miniprep Kit (Qiagen) and screened for the presence of T375A by Sanger sequencing. Susceptibility to DRV, atazanavir, LPV, indinavir and saquinavir, and replication capacity were determined by a single-cycle assay in HEK293T cells as described.^19^ Virus replication in the presence of the drug was determined by luciferase quantification 48 hours post-infection relative to no-drug controls. The mean 50% inhibitory concentration (IC50) from 3 separate experiments was calculated and results expressed as fold-changes (FC) in IC50 compared to wild-type P8.9NSX. Viral replicative capacity was determined by luciferase quantification in the absence of drug, calculating the mean luciferase activity from ≥4 values within the linear range, and expressed as percentage relative to wild-type P8.9NSX.

## Results

### Study population

At study entry, the 120 participants had received ART for a median of 7.5 years, including prior first-line NNRTI-based ART for a median of 3.0 years and current second-line PI/r-based ART for a median of 3.1 years (Table 1). Most (106/120, 88.3%) were receiving once daily TDF/3TC plus twice daily LPV/r. The characteristics of the two arms were overall similar, although the DRV/r arm showed higher levels of total HIV-1 DNA and higher prevalence of RAMs in HIV-1 DNA (Table 1). As per the eligibility criteria, prior to randomisation all participants had shown a viral load <60 copies/ml in 2 screening measurements, which were separated by a median of 7 weeks (range 6-12). No prior viral load measurements were available in the medical records.

### Resistance patterns in cellular HIV-1 DNA at study entry

HIV-1 DNA sequences were obtained from 90/120 (75%) participants (Table 1). Six participants did not have a PBMC sample and 24 did not yield a sequence in ≥2 attempts (until sample exhaustion). Overall 58/90 (64.4%) sequences showed ≥1 RAM, most commonly affecting the NRTIs (52/90, 57.8%) and the NNRTIs (53/90, 58.9%); 47/90 (52.2%) had RAMs for both classes (Table 1). Protease RAMs were uncommon; 3 participants showed the nelfinavir RAM D30N either alone or with RAMs to other classes; they had received LPV/r for 1-3 years and had no prior nelfinavir exposure. Overall, 11/90 (12.2%) HIV-1 DNA sequences showed in-frame stop codons within RT, including 9 sequences containing RAMs; hypermutation was also common in both RT and protease sequences containing RAMs (Table 2).

**Table 2.**
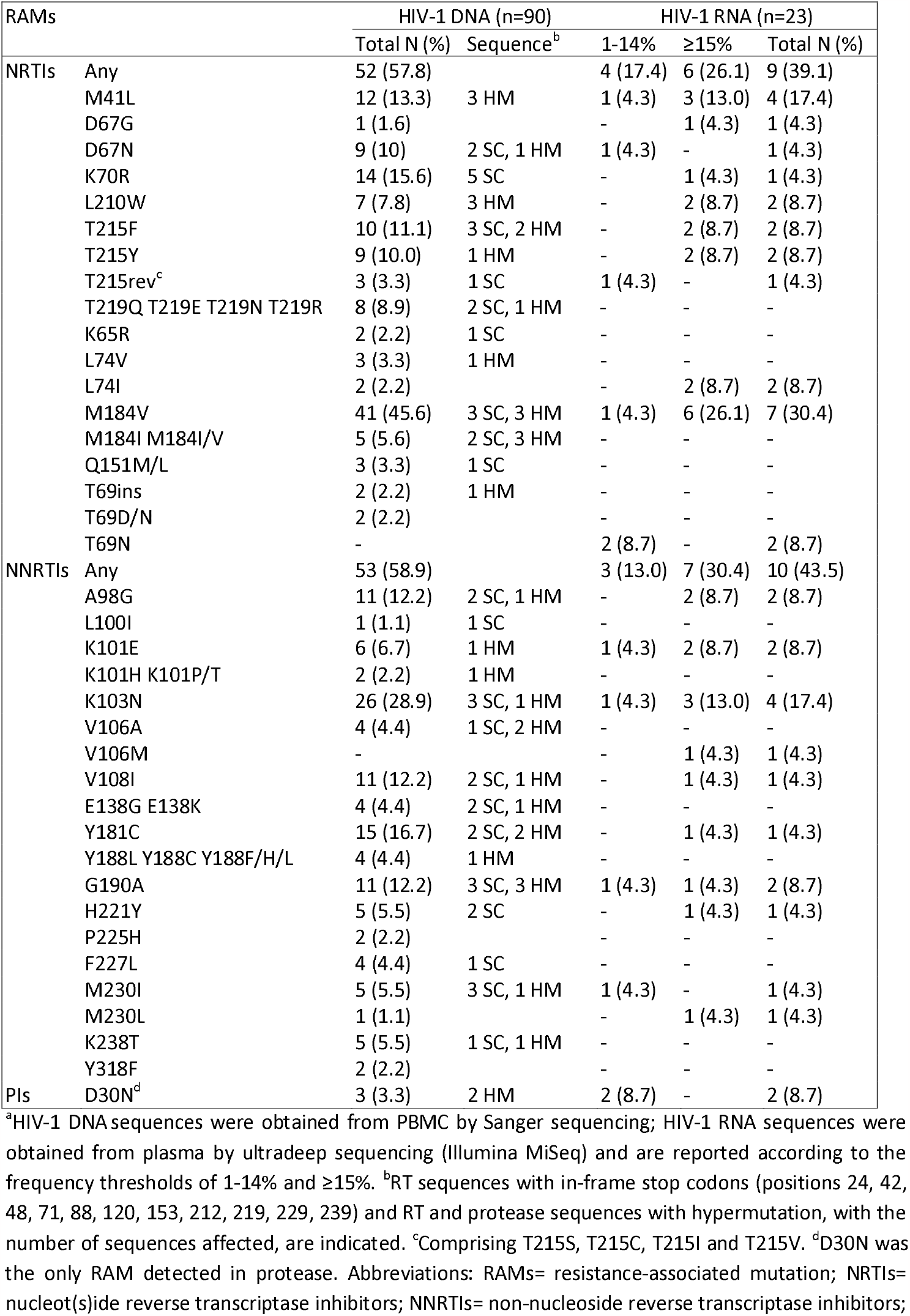
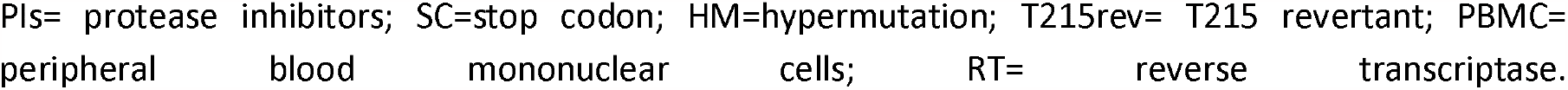
Major resistance-associated mutations in cellular HIV-1 DNA at study entry (n=90) and in plasma HIV-1 RNA at virological rebound (n=23)^a^.

### Virological outcomes

At week 24, proportions with HIV-1 RNA <400 copies/ml (primary endpoint) were 72/81 (88.9%) in the DRV/r arm and 37/39 (94.9%) in the SOC arm (p=0.50). Proportions with HIV RNA <60 copies/ml through week 24 were 62/81 (76.5%) and 36/39 (92.3%), respectively (p=0.04). In the DRV/r arm, 24/81 (29.6%) participants experienced virological rebound (confirmed or last available HIV-1 RNA ≥60 copies/ml) through week 48, including 16/81 (19.8%) who experienced virological failure (confirmed or last available HIV-1 RNA ≥400 copies/ml). In the SOC arm, 3/39 (7.7%) participants experienced virological rebound through week 24, including 1/39 (2.6%) participant with virological failure.

### Resistance patterns in plasma HIV-1 RNA at virological rebound

Sequences were obtained by UDS in 23 participants who experienced virological rebound (DRV/r arm n=21; SOC arm n=2). HIV-1 RNA levels at the time of testing were median 3.0 log10 copies/ml (range 2.0-4.1). NRTI and NNRTI RAMs were found in 9/23 (39.1%) and 10/23 (43.5%) samples, respectively and 8/23 (34.7%) samples had RAMs for both classes; most RAMs occurred at frequency ≥15% (Table 2). No participant had DRV RAMs at either high or low frequency. Two participants, both in the DRV/r arm, showed the nelfinavir RAM D30N at low frequency (3-4%); before starting DRV/r, they had received LPV/r for >4 years and had no prior nelfinavir exposure. No stop codons or hypermutation were found in plasma sequences.

### Comparison of resistance patterns in HIV-1 DNA and HIV-1 RNA

There were 18 participants with paired cellular HIV-1 DNA and plasma HIV-1 RNA sequences (Table 3). Phylogenetic analyses confirmed clustering of their *pol* sequences (Figure 1). With 6/18 (33.3%) sample pairs, the resistance patterns were fully consistent, comprising 5 with no RAMs in either sample and 1 with the NNRTI RAM K103N in both samples (Table 3). A further 6/18 (33.3%) sample pairs showed partial consistency, comprising 5 with fewer RAMs in HIV-1 DNA and 1 with fewer RAMs in HIV-1 RNA. With 5/18 (27.8%) sample pairs, RAMs were detected only in HIV-1 DNA; 3 HIV-1 DNA sequences showed in-frame stop codons and 1 with the NNRTI RAM M230I also showed hypermutation. One sample pair showed M230I in HIV-1 RNA only (frequency 1%).There was no consistency in the detection of the protease RAM D30N when comparing sample pairs, and no linkage was detected between *pol* sequences from the 5 individuals with D30N (Supplementary Figure 1).

**Table 3.**
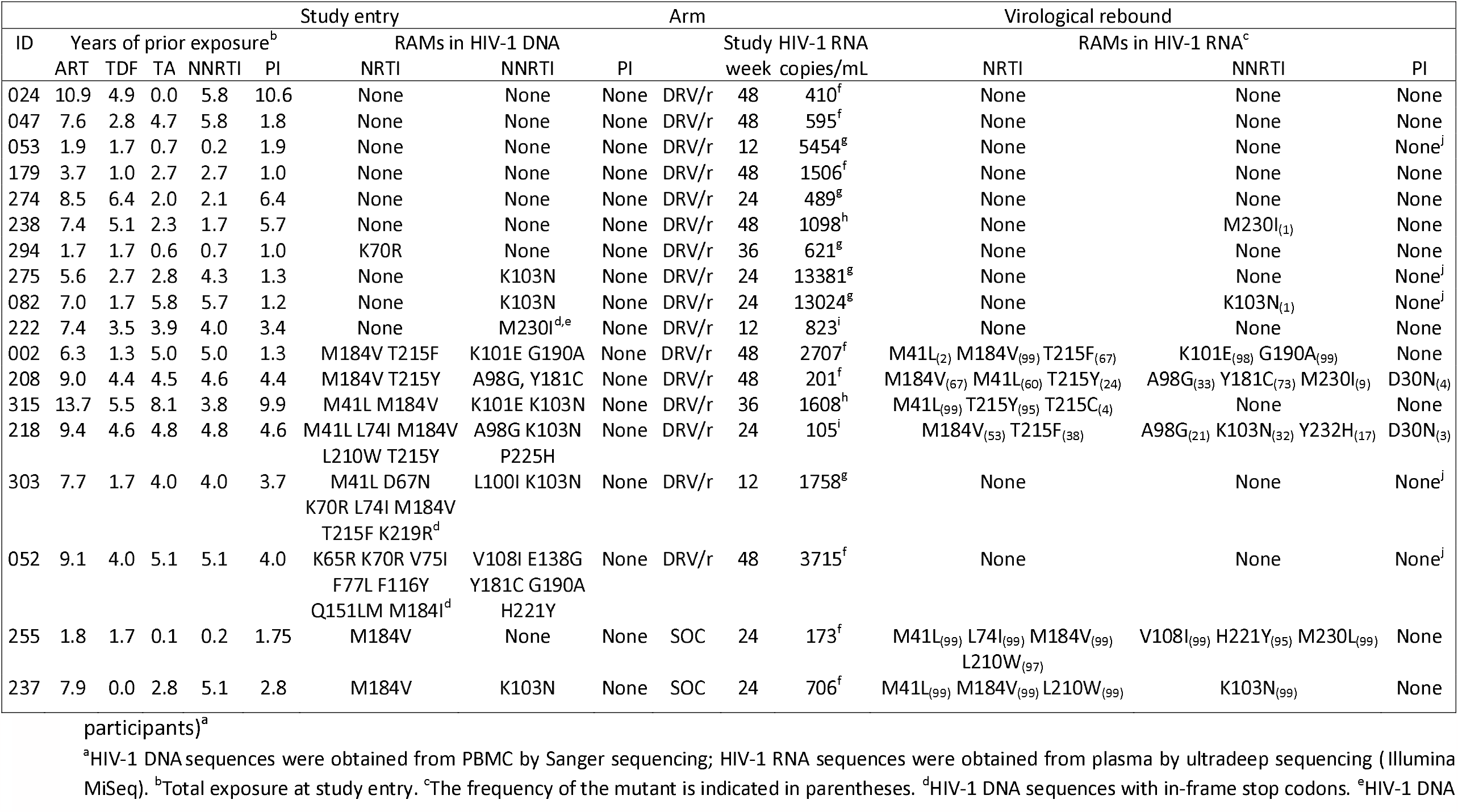
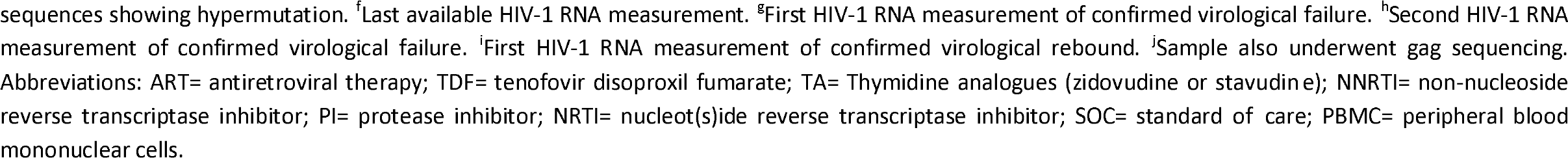
Paired cellular HIV-1 DNA sequences obtained at study entry and plasma HIV-1 RNA sequences obtained at virological rebound (n=18 participants)^a^.

**Figure 1.**
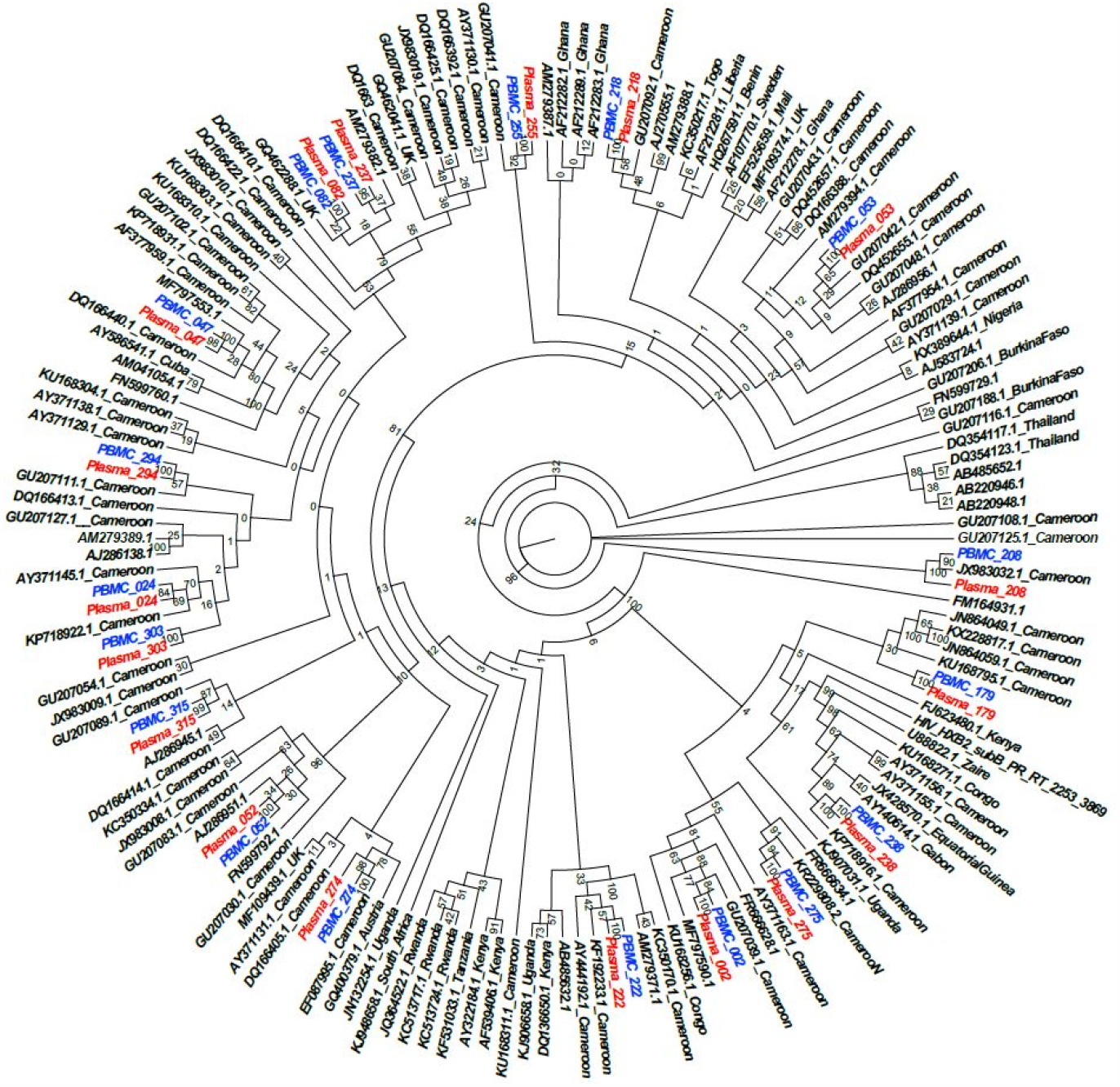
Maximum-likelihood phylogenetic tree of paired *pol* sequences of HIV-1 DNA obtained from PBMC at study entry (blue) and HIV-1 RNA obtained from plasma at virological rebound (red) (n=18 pairs). Control sequences were obtained from Genbank (duplicate sequences excluded). The tree was inferred using 1000 bootstrap replicates.

### Gag mutations

Among participants who experienced virological rebound on DRV/r, 8 underwent gag sequencing, including 4 who were tested longitudinally (Table 4). None of the sequences contained RAMs in protease. Screening for gag mutations significantly associated with PI exposure revealed 4 sequences with 5 cleavage site mutations (K436R,. One participant with longitudinal samples showed emergence of K436R in p7/p1 between week 24 and week 36, when K436R replaced T375N in p2/p7. Two sequences showed T375A in p2/p7, in one case occurring with K436R. The phenotypic effects of T375A were analysed by site-directed mutagenesis. The mutation reduced PI susceptibility from around 5-FC for atazanavir, indinavir and LPV to 10-FC for DRV and saquinavir (Figure 2). The mutation also increased replicative capacity to 161% (±5.0) relative to wild-type control. furthermore, all sequences showed mutations within other gag regions. Among participants with longitudinal results, emergent mutations included D121G in p17, T190I in p24 and P478Q in p6.

**Table 4.**
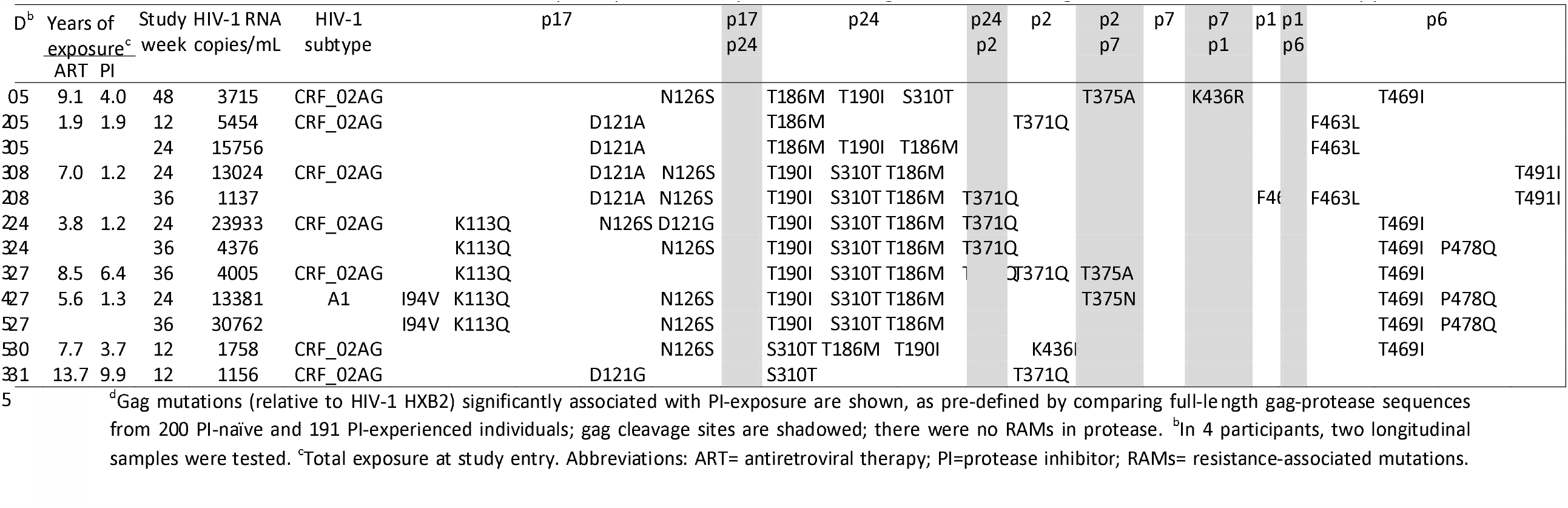
Gag mutations in plasma HIV-1 RNA of participants who experienced virological rebound during darunavir/ritonavir monotherapy^a^.

**Figure 2.**
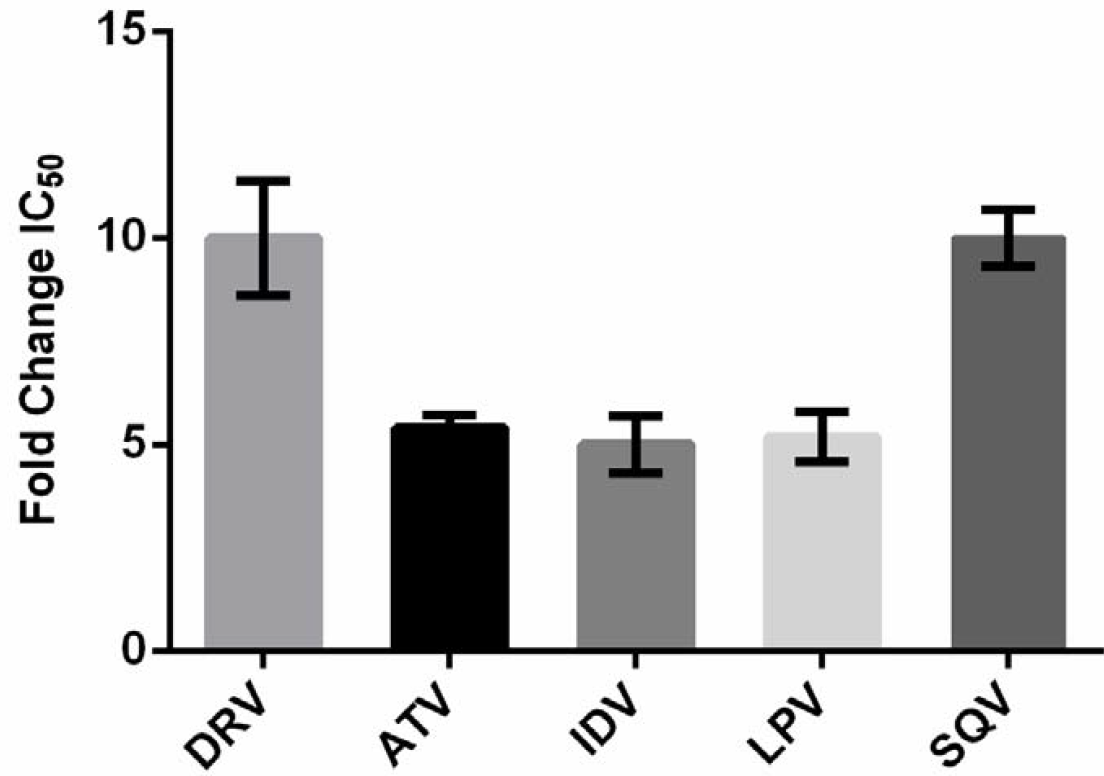
Phenotypic resistance profile of the gag p2/p7 cleavage site mutation T375A, as determined by site-directed mutagenesis. The mean fold-change in IC50 from 3 independent experiments (with standard deviation) is shown. Abbreviations: ATV= atazanavir; DRV= darunavir; IDV= indinavir; LPV= lopinavir; SQV= saquinavir.

## Discussion

We studied a population in Cameroon that following failure of first-line NNRTI-based ART started 2 NRTIs plus a PI/r (typically LPV/r) in the absence of virological monitoring. After confirmation of virological suppression, participants were assigned to either maintenance monotherapy with DRV/r or TDF/3TC plus LPV/r. The findings align with published data from a similar population,^6^ indicating a heightened risk of viremia among individuals on DRV/r monotherapy. Over 48 weeks, we observed several instances of virological rebound (confirmed or last available HIV-1 RNA ≥60 copies/ml) and virological failure (HIV-1 RNA ≥400 copies/ml) in this group, with nearly 30% of participants experiencing rebound viremia. At virological rebound, the resistance patterns in plasma HIV-1 RNA were partially reflective of those detected in cellular HIV-1 DNA at study entry. UDS confirmed the absence of treatment-emergent DRV RAMs in protease. However, we detected mutations in gag and demonstrated an effect on DRV susceptibility in the absence of RAMs in protease.

In cell culture, two pathways of DRV resistance have been characterised anchored around the protease RAMs I50V or I84V.^20^ DRV RAMs have also been shown to emerge in PI-experienced people with pre-existing protease RAMs but have rarely occurred when starting DRV/r de novo; the high resistance barrier is thought to reflect tight binding of DRV to the protease enzyme and high plasma concentrations achieved through pharmacological boosting.^20^ Monotherapy studies have also reported a negligible risk of DRV RAMs.^5^ Most studies employed Sanger sequencing for detecting resistance, but 2 studies applied more sensitive techniques. One study using single genome sequencing in 5 participants with HIV-1 RNA >400 copies/ml detected DRV RAMs (V32I, I47V, I50V) in one participant lacking the mutations by Sanger sequencing.^21^ A second study using UDS in 14 participants with HIV-1 RNA >1000 copies/ml, found I54T in the protease of a participant lacking protease RAMs by Sanger sequencing;^22^ however, I54T is not considered a DRV RAM.^11^ We complement these data by reporting UDS results from 21 participants and by extending the analysis to all cases of confirmed or last available plasma HIV-1 RNA ≥60 copies/ml, including 12 with viral load <1000 copies/ml. No DRV RAMs were detected in the 21 participants.

Two participants on DRV/r showed the protease RAM D30N in rebound HIV-1 RNA but not in HIV-1 DNA at study entry. The significance is doubtful. D30N is a non-polymorphic substrate-cleft mutation that is selected by and causes high-level resistance to nelfinavir.^23^ The aspartate-to-asparagine substitution alters the hydrogen bond interaction with the aniline NH2 group of DRV causing some loss of binding affinity.^24^ However, the mutation occurred only at low frequency (<5%) in rebound HIV-1 RNA and there is no evidence of an association with DRV resistance.^11^ At study entry, D30N was also detected in the HIV-1 DNA of 3 LPV/r-experienced participants, but there is similarly no evidence that LPV/r selects for D30N.^23^ PCR-induced error or APOBEC3G-mediated hypermutation, an innate defence mechanism that aims to impar virus functionality, may also explain the unexpected detection of D30N.^25^ Substitutions that have been related to hypermutation include D30N and M46I in protease and E138K, M184I, G190E and M230I in RT.^25-27^ We found hypermutation in several HIV-1 DNA sequences showing these mutations.

Mutations in gag, the natural substrate of the protease enzyme, can improve gag-protease binding and gag processivity and modulate PI susceptibility and viral fitness in the absence or alongside protease RAMs.^28-34^ Mutations associated with PI exposure typically involve gag cleavage sites, although mutations in other regions have also been implicated.^18,33,34^ We had sufficient sample for full-length gag and protease sequencing in a subset of participants experiencing virological rebound on DRV/r. All had gag mutations previously associated with PI exposure, including mutations at the cleavage sites p2/p7 and p7/p1 and within p17, p24, p2 and p6. In one case, we were able to demonstrate emergence of K436R in p7/p1 while HIV-1 RNA levels ranged between 13,381 and 30,762 copies/ml on DRV/r. K436R was associated with PI exposure and resistance in previous studies and shown to occur during DRV/r monotherapy.^28,33^ We also detected T375A in p2/p7 in 2 participants. T375A was previously detected in an individual receiving DRV/r monotherapy in combination with S451N in p1/p6 and several additional gag mutations.^33^ Given the limited data available, we produced a site-directed mutant with T375A and observed 10-FC reduction in DRV susceptibility, alongside increased viral replication capacity. Outside of cleavage sites, we saw emergence of D121G in p17, T190I in p24 and P478Q in p6. T190I is an HLA-B*81-associated mutation with a compensatory role in viral fitness.^35^ Regrettably, we were unable to expand gag analyses due limited sample volumes. Of note, we focused the analysis of gag sequences on mutations previously found to be associated with PI-exposure, using strict criteria to define the association. This approach may not account for all possible gag mutations that may contribute to DRV/r resistance. In fact, significant mutations may not necessarily be confined to consistent changes at a few sites.^36^ Importantly, all participants with gag mutations regained virological suppression after receiving adherence support and returning to PI/r-based triple ART.

There is growing interest in the potential clinical utility of sequencing cellular HIV-1 DNA to guide treatment decisions during virological suppression.^2^ As we previously reported on a smaller subset,^4^ the resistance patterns detected in HIV-1 DNA at study entry were consistent with the previous failure of NNRTI-based ART. The resistance patterns detected in rebound HIV-1 RNA on DRV/r were generally in agreement with those found at study entry, but discrepancies were common. This is not unexpected. Several HIV-1 DNA sequences containing RAMs were defective and could not be expected to sustain virus production.^37^ A further consideration is that in the setting of virological suppression the HIV-1 DNA input into the sequencing test is small and detection of RAMs can become stochastic. Due to limited infrastructure, we did not perform UDS on PBMC, which may have increased the detection of RAMs in HIV-1 DNA,^38^ although even UDS cannot be expected to overcome issues of input size and provide a full representation of the resistance archive.^2^

In summary, the resistance patterns of cellular HIV-1 DNA sequenced during virological suppression, whilst largely reflective of treatment history, are only partially consistent with the resistance patterns of rebound viremia. Both biological and technical factors may account for the discrepancies and should be taken into consideration when applying the test clinically. We confirm the high resistance barrier of DRV/r but demonstrate that gag substitutions, including T375A in in p2/p7, provide an alternative pathway of DRV resistance.

## Acknowledgments

We are grateful to the study participants and the staff of Hôpital Central Yaoundé and the Centre Pasteur of Cameroon.

## Transparency declarations

AMG reports personal payments from Abbott, Gilead, GSK, Roche and ViiV, and grants to the institution from Gilead and ViiV, outside of the present work.

